# Ancient RNA virus epidemics through the lens of recent adaptation in human genomes

**DOI:** 10.1101/2020.03.18.997346

**Authors:** David Enard, Dmitri A. Petrov

## Abstract

Over the course of the last several million years of evolution, humans likely have been plagued by hundreds or perhaps thousands of epidemics. Little is known about such ancient epidemics and a deep evolutionary perspective on current pathogenic threats is lacking. The study of past epidemics has typically been limited in temporal scope to recorded history, and in physical scope to pathogens that left sufficient DNA behind, such as *Yersinia pestis* during the Great Plague.

Host genomes however offer an indirect way to detect ancient epidemics beyond the current temporal and physical limits. Arms races with pathogens have shaped the genomes of the hosts by driving a large number of adaptations at many genes, and these signals can be used to detect and further characterize ancient epidemics.

Here, we detect the genomic footprints left by ancient viral epidemics that took place in the past ~50,000 years in the 26 human populations represented in the 1,000 Genomes Project. By using the enrichment in signals of adaptation at ~4,500 host loci that interact with specific types of viruses, we provide evidence that RNA viruses have driven a particularly large number of adaptive events across diverse human populations. These results suggest that different types of viruses may have exerted different selective pressures during human evolution. Knowledge of these past selective pressures will provide a deeper evolutionary perspective on current pathogenic threats.

## Introduction

About 40 years ago and for the first time in their evolution, humans eradicated a virus that had claimed countless lives, the Variola virus known as the causal agent of smallpox (Metzger et al., 2015). Since then, and despite progress in prevention, many new viral threats have emerged and spread, including viruses such as HIV (Hemelaar, 2012; Worobey et al., 2008), Ebola virus (Breman et al., 2016), Zika virus (Gubler et al., 2017), and SARS coronavirus (Holmes and Rambaut, 2004). The number of new viral zoonoses (transmission from animals to humans) keeps increasing as a result of multiple factors notably including higher population density and the disturbance of wild habitats (Jones et al., 2013). Although the current frequency of zoonoses may be unusually high due to large human populations in contact with other species, even much less frequent viral zoonoses during the millions of years of past human evolution are likely to have resulted in many epidemics.

When viruses impose a selective pressure on a host, adaptation happens in response at the host loci that interact with the virus (Enard et al., 2016). An epidemic can then be detected through the enrichment in signals of adaptation at VIPs that interact with a virus. One obstacle to utilizing this method is that we do not have a direct access to the interactions between a host and an ancient virus. However, host-virus interactions are well known in present viruses. Because phylogenetically related pathogens tend to use the same interactions with their hosts (Enard et al., 2016), we can use present interactions as proxies for the interactions with ancient viruses from the same phylogenetic family. For example, HIV belongs to the family of lentiviruses, and we can use interactions between HIV and human proteins as proxies for ancient interactions between human proteins and ancient lentiviruses.

Accordingly, we recently found that arms races with ancient viruses have left characteristic signals of abundant, intense adaptation in human proteins that interact with present viruses (Castellano, 2019; Enard et al., 2016; Uricchio et al., 2019). According to our curation and annotation of the virology literature, in the past three decades, virologists have found that approximately 4,500, or 20% of human proteins physically interact with human-infecting viruses (Enard and Petrov, 2018). A large proportion of these interactions have known functional consequences for the viral replication cycle, thus making virus-interacting proteins (VIPs) prime candidates for host adaptation in response to viral epidemics (Methods and Table S1). VIPs harbor remarkably high levels of past protein adaptation, with rates of adaptive amino acid changes several times higher than human proteins that are not known to interact with viruses (non-VIPs) (Enard et al., 2016; Uricchio et al., 2019). Furthermore, adaptation was not only more frequent at VIPs compared to non-VIPs; it was also stronger, more intense adaptation (Castellano, 2019; Uricchio et al., 2019), suggesting that viruses repeatedly imposed strong selective pressures on their human hosts during evolution. Thus, frequent new zoonoses and abundant adaptation at VIPs together suggest that viruses drove many epidemics in past human evolution.

Despite these clear indications that ancient viral epidemics may have been frequent, little is known about which ancient viruses were involved, and how ancient viruses that caused ancient epidemics relate phylogenetically to current viruses infecting humans. The current lack of knowledge of ancient viruses can be explained by the difficulty to recover ancient viral DNA (Spyrou et al., 2019), and by the fact that many viruses have their genomes coded by RNA that is known to degrade much faster than DNA. In light of the scarcity of molecular remains, how can we identify the viruses that drove ancient epidemics during human evolution?

In order to identify viruses that drove ancient epidemics, we can use present interactions as proxies for the interactions with ancient viruses from the same phylogenetic family. Using this approach, we previously found evidence that 50,000 years ago (Abi-Rached et al., 2011; Fu et al., 2015; Green et al., 2010; Paabo, 2015; Sankararaman et al., 2014), Neanderthals appear to have infected the modern human ancestors of present Europeans with one or multiple RNA viruses, as shown by the fact that European modern humans harbor substantially more and substantially longer introgressed Neanderthal DNA at genes that interact with RNA viruses compared to genes that interact with DNA viruses (Enard and Petrov, 2018). The key factor in this finding was that ancient RNA viruses drove adaptive introgression of Neanderthal DNA not only at a few loci, but at dozens of loci, thus creating a strongly significant enrichment of Neanderthal DNA at RNA VIPs (VIPs that interact with RNA viruses) compared to DNA VIPs (VIPs that interact with DNA viruses). The magnitude of the adaptive signals is what enabled their assignment to a specific type of viruses, in this case RNA viruses.

Because when they interbred (Abi-Rached et al., 2011; Fu et al., 2015; Green et al., 2010; Paabo, 2015; Sankararaman et al., 2014), Neanderthals and modern humans likely infected each other with their respective viruses, there was a strong prior to expect ancient epidemics at the time of interbreeding, and it made sense to start looking for host genomic signals of ancient epidemics at this particular time of human evolution. However, because ancient epidemics were likely frequent during human evolution, it now also makes sense to extend the search for signals of adaptation left by ancient epidemics to any time of human evolution from which we can get signals of adaptation. Recent human adaptation is particularly interesting in this respect, because we now have good genome datasets (Genomes Project et al., 2015) and statistical tools (Ferrer-Admetlla et al., 2014; Schrider and Kern, 2016, 2017, 2018; Voight et al., 2006) to detect signals of recent adaptation genome-wide in the form of recent selective sweeps. Haplotype-based statistics that use the structure of haplotypes along chromosomes have been shown to be particularly useful to detect recent selective sweeps, because they have good statistical power to detect strong, recent incomplete sweeps (Ferrer-Admetlla et al., 2014; Garud et al., 2015; Voight et al., 2006) without suffering from the confounding effect of other processes such as background selection (Enard et al., 2014). Among available haplotype-based statistics, the iHS or integrated Haplotype Score (Voight et al., 2006) also has the three advantages, (i) of having been extensively tested (Barreiro et al., 2009; Enard et al., 2014; Johnson and Voight, 2018; Voight et al., 2006), (ii) of showing versatility when it comes to detect sweeps from de novo mutations or from standing genetic variation (see power analysis in Methods), and (iii) of having fast implementations that can be used to scan many individual genomes from many human populations in a non-prohibitive amount of time (Maclean et al., 2015) (Methods).

Here, we examine signals of recent human adaptation in diverse human populations to ask whether viruses drove an enrichment of signals of recent adaptation at VIPs compared to non-VIPs. Specifically, we test whether VIPs are overall enriched for recent selective sweeps detected by the well-established iHS statistic in the 26 human populations represented in the 1,000 Genomes phase 3 dataset (Genomes Project et al., 2015). The iHS statistic has better power to detect the most recent, incomplete selective sweeps, and no power to detect selective sweeps older than 50,000 years (see power analysis, Methods). This restricts our analysis of recent adaptation in response to viruses to human evolution after the migration out of Africa.

We further ask whether specific types of viruses drove recent adaptation more frequently than others. More specifically, our previous results on adaptive introgression from Neanderthals to modern humans (Enard and Petrov, 2018), and the known zoonoses from the recent past (Geoghegan et al., 2017; Kreuder Johnson et al., 2015), suggest that RNA viruses may have driven more recent adaptation than DNA viruses because they jump more often from a species to another. The vast majority of known zoonotic viruses that infected humans in the recent past are RNA viruses from diverse RNA virus phylogenetic families, ranging from lentiviruses such as HIV, flaviviruses (Dengue and Zika virus), filoviruses (Ebola virus), to orthomyxoviruses (Influenza virus). Comparatively, most DNA viruses infecting humans were transmitted from animals much longer ago (Geoghegan et al., 2017) and thus might cross species barriers less frequently in general. DNA viruses also tend to include less pathogenic viral families such as herpesviruses (Petti and Lodi, 2019).

Taken together, these diverse lines of evidence suggest the hypothesis that RNA viruses might have imposed a stronger selective pressure than DNA viruses during recent human evolution. To test this, we compare the enrichment in iHS selective sweeps at RNA VIPs and the same enrichment at DNA VIPs. The dataset of 4,500 VIPs is particularly well suited for this comparison, with similar numbers of RNA and DNA VIPs (2,691 and 2,604, respectively, Table S1), providing an evenly powered comparison.

First, using all VIPs compared to non-VIPs, we find a strong enrichment of iHS selective sweeps at VIPs overall, suggesting that viruses were a major selective pressure during recent human evolution that drove multiple strong adaptive events. Second, we further find that the selective sweeps enrichment is much more pronounced at RNA VIPs compared to DNA VIPs. These results validate previous results and are consistent with RNA viruses being a significant selective pressure in the past 50,000 years of human evolution.

## Results

### Properties of VIPs

We use a dataset of ~4,500 VIPs that are human proteins known to interact physically with viruses (Table S1). Of these, we previously manually curated 1,920 from the virology literature (Methods). The articles that report these interactions also frequently report their effect on the viral replication cycle, and 66% of the manually curated interactions also have reported proviral (interaction is beneficial to viral proliferation), or antiviral effects (interaction is detrimental to the virus) (Methods). While this percentage is high, it is still likely to be an underestimate due to the fact that it is limited to the cases where the effect of the interaction has been investigated in the first place. This makes VIPs prime candidates for host adaptation, with a strong likelihood of functional mutations affecting the viral replication cycle. The remaining ~2,600 VIPs were identified using high-throughput methods and were retrieved from the VirHostnet 2.0 database as well as from a number of additional studies (Table S1). For this analysis, we pooled the manually curated and the high-throughput VIPs together into one group that we systematically compared to the rest of the genome. In total, 20 viruses that infect humans have more than 10 VIPs and 14 viruses have more than 100 VIPs, with influenza virus (IAV) and HIV having the highest numbers of known VIPs (1,505 and 1,209 respectively; Table S1). Note that VIPs and all other genes used for this analysis are Ensembl v83 genes (Zerbino et al., 2018).

A simplifying property of this set of VIPs for our analysis is that they are almost exactly evenly distributed between VIPs that interact with RNA viruses (2,691 RNA VIPs) and VIPs that interact with DNA viruses (2,604 DNA VIPs), with 1,134 VIPs interacting with both. This similar number of interactions makes it possible to compare sweeps at RNA VIPs with sweeps at DNA VIPs with no bias in statistical power.

### VIPs are enriched for recent selective sweeps in human populations

We first test if VIPs are overall enriched for recent selective sweeps compared to non-VIPs. We use the iHS statistic to detect candidate sweeps in each of the 26 populations from the 1,000 Genomes Project (Maclean et al., 2015) (Methods). The iHS statistic is measured for a focal variant (Voight et al., 2006), by measuring how far the haplotypes carrying the derived allele of the focal variant extend both upstream and downstream of it, compared to how far the haplotypes carrying the ancestral allele extend. A rapid increase in frequency of the derived focal allele due to strong positive selection results in large and frequent linked haplotypes compared to the haplotypes carrying the ancestral allele, and in elevated values of the iHS statistic. We first measure iHS along chromosomes for all variants in all the 26 populations (Methods), in a way that optimizes the sensitivity of iHS to strong selective sweeps (see power analysis in Methods). We do this because our previous work using a very different methodology to quantify adaptation (the McDonald-Kreitman test) (McDonald and Kreitman, 1991; Tataru et al., 2017; Uricchio et al., 2018) estimated that adaptation at VIPs was not only frequent but also generally driven by strongly adaptive mutations (Uricchio et al., 2019), which predicts strong selective sweeps at VIPs.

We then rank all protein coding genes in the genome from highest iHS values to lowest iHS values. We measure the average iHS in large 1,000kb windows centered on the genomic center of genes. These large windows are more specifically sensitive to strong adaptation compared to smaller windows, and using a constant window size avoids biases due to gene length (Methods). For example, the top 200 genes are then the 200 genes in a population with the highest iHS compared to other genes, the top 1,000 are the 1,000 genes with the highest iHS, and so on. Finally, we count how many VIPs are among the top-ranking genes in a specific population. We then sum the number of VIPs among top-ranking genes across all the 26 populations. That is, if the world included three populations A, B and C, and the respective top 100 of populations A, B and C include 23, 34, and 15 VIPs, then the worldwide top 100 would be 23+34+15=72. We then measure the same sum but in the control sets of non-VIPs, and compute the corresponding fold enrichment in VIPs compared to the average of the control sets of non-VIPs. This implies that the results represent average worldwide trends.

Importantly, the sets of control non-VIPs account for multiple key potential confounding factors (Methods). Indeed, VIPs and non-VIPs not only differ by the fact that VIPs are not to interact with viruses while non-VIPs either do not interact with viruses or are not known to. In addition, VIPs and non-VIPs differ by many other factors (Enard et al., 2016; Enard and Petrov, 2018). For instance, VIPs are more highly constrained, more highly expressed than non-VIPs, and have more protein-protein interactions than non-VIPs (Enard et al., 2016; Enard and Petrov, 2018). If gene expression, protein-protein interactions (Luisi et al., 2015), or other factors affect the prevalence of recent sweeps on their own, independently of interactions with viruses, they might confound the comparison of VIPs and non-VIPs by creating differences between the former and the latter that have nothing to do with interactions with viruses. Thus, we build random sets of control non-VIPs that match VIPs for multiple potential confounding factors using a previously described bootstrap test (Castellano, 2019) (Methods). Furthermore, the control sets of non-VIPs exclude all non-VIPs that are too close to VIPs and may thus be found in the same large sweeps extending over multiple genes. To avoid counting large VIP sweeps also as non-VIP sweeps, or large non-VIP sweeps also as VIP sweeps, we select only control non-VIPs at least 500kb away from VIPs. The enrichment of recent adaptation is therefore tested comparing sweep signals at VIPs compared to sweeps signals far from VIPs. In addition, to avoid the confounding effect of counting the same adaptive event multiple times when VIPs are clustered together in the same selective sweep, we use an approach based on block-randomized genomes to assess the statistical significance of the signals detected (Methods).

We retrieved the number of control non-VIPs among the top-ranking sweep genes for each of the 1,000 control sets, measured the corresponding average, and measured how enriched VIPs were in top-ranking genes compared to this control average. Furthermore, we measured the fold enrichment at VIPs for several sets of iHS top-ranking genes corresponding to different rank thresholds. The top-ranking genes are defined by sliding a rank threshold from the top 2,000 genes, to a much more stringent threshold for the top 20 genes (see figure 1, x axis). The top 2,000 genes include even weak sweep signals, while the top 20 genes only correspond to strong sweep candidates. Using a sliding rank threshold thus avoids making assumptions on exactly how strong adaptation to viruses needs to be, and enables the detection of enrichments for strong selective sweeps (top 100 or less) as well as enrichments driven by more incomplete or weaker, polygenic hitchhiking signals spread across more genes (see power analysis in Methods).

**Figure 1.**
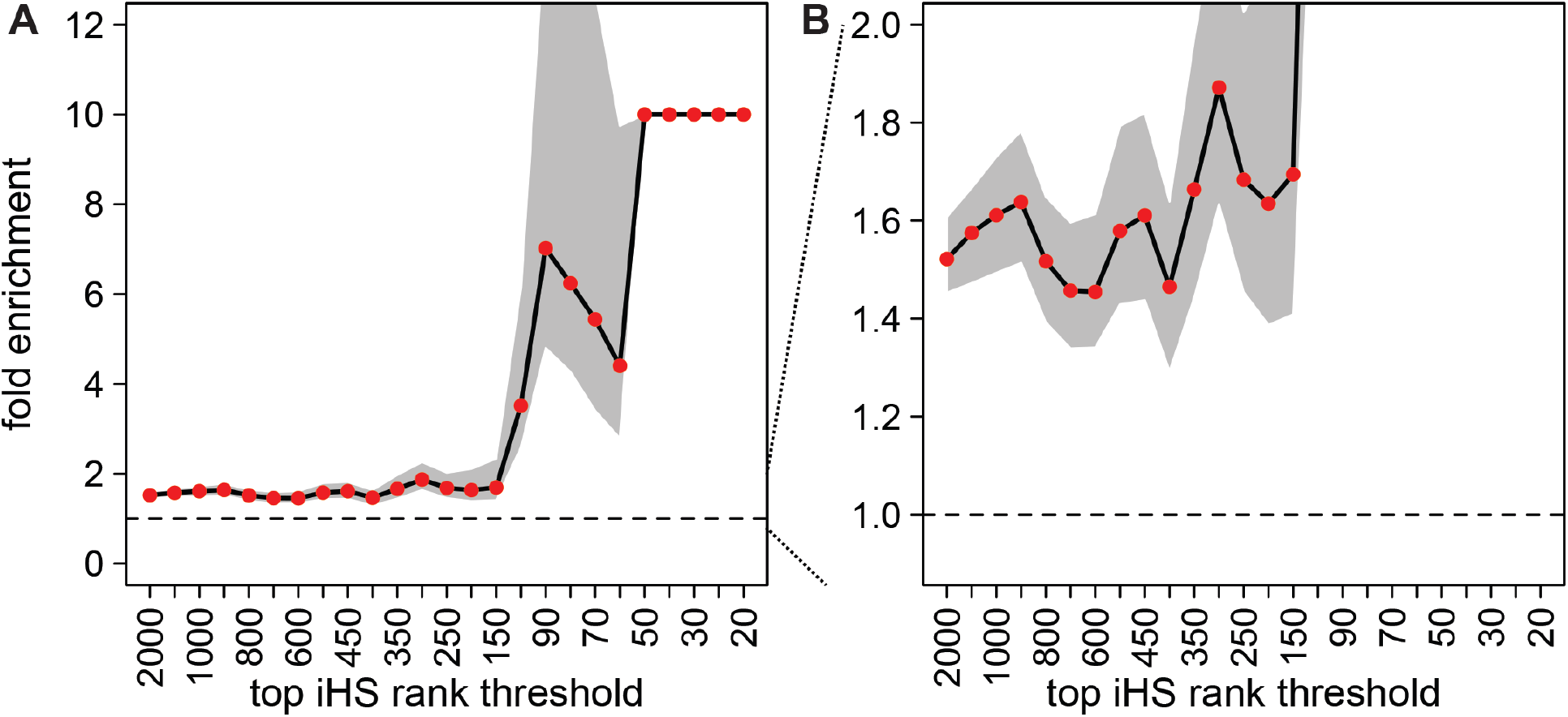
Worldwide enrichment of iHS sweeps at VIPs compared to control non-VIPs. A. Black line: observed fold enrichment at VIPs. Grey area: 95% confidence interval of the fold enrichment. Fold enrichments above ten are represented at ten. When this happens, the confidence interval is not represented. However the lowest edge of the confidence intervals not represented are all above one. Red dots: bootstrap test P<0.001 (Methods). Dashed line: fold enrichment of one, i.e. no enrichment. Fold enrichment (y-axis) is the number of VIPs in candidate sweeps divided by the average number of control-non-VIPs in candidate sweeps. VIPs and non-VIPs in candidate sweeps are counted if they belong to the top x iHS genes (x-axis), where x is a rank threshold that slides from top 2,000 to top 20, taking in total 25 values (2,000; 1,500; 1,000; 900; 800; 700; 600; 500; 450; 400; 350; 300; 250; 200; 150; 100; 90; 80; 70; 60; 50; 40; 30; 25; 20). A fold enrichment of y=3.51 at top x=100 means that there are 3.51 times more VIPs in the top 100 iHS genes than control non-VIPs on average (over 1,000 control sets of non-VIPs). There are in fact 80 VIPs in the iHS top 100, versus only 22.7 control-non-VIPs. 80 is high compared to 100 because of the summing over all 26 human populations. Specifically, a VIP or non-VIP counts as one in the top x if it is in the top x of at least one of the 26 populations. Note that counting the number of genes instead of counting the number of sweeps ignores the clustering of multiple genes in a single sweep, but that we account for this potential bias when estimating the whole rank threshold curve significance (see Methods).

Using this strategy, we find that VIPs are enriched for recent selected sweeps with confidence intervals clearly above the no-enrichment mark (fold enrichment of one in figure 1A,B, dashed horizontal line). VIPs are enriched for genes from the top 2,000 of iHS (figure 1A,B), suggesting an excess of weak to moderate hitchhiking events. The relative enrichment shown in figure 1 at top 2,000 corresponds to an absolute number of 527 additional VIPs compared to control expectations. VIPs are however particularly enriched for strong selective sweep signals in the top 100 of iHS or less (figure 1A), suggesting that viruses drove particularly strong adaptive events in recent human evolution. The iHS to 100 includes 80 VIPs in sweeps in different human populations, versus only 22.7 expected by chance according to the bootstrap test. The enrichment observed for the top 100 thus cannot explain alone the enrichment observed for the top 2,000 (57 vs. 527 additional VIPs, respectively). Overall, the enrichments from the top 2,000 to the top 20 are collectively highly significant and robust to issues such as clustering of multiple VIPs in the same sweep (P<0.001; see Methods describing how we estimate the significance of the whole enrichment curve and not just at one rank threshold). These results suggest that viruses may have driven multiple strong adaptive events during that past ~50,000 years of human evolution, and are consistent with our previous results showing a high enrichment of adaptive events at VIPs at different evolutionary time scales (Enard et al., 2016; Enard and Petrov, 2018; Uricchio et al., 2019).

### RNA viruses drove more recent selective sweeps than DNA viruses

The strong excess of iHS selective sweeps at VIPs suggests that there may be enough statistical power to cut the set of VIPs further into smaller categories, and ask which types of viruses drove this signal. Specific viruses may have indeed driven more epidemics than others in recent evolution, and a completely homogeneous distribution of sweeps between the VIPs of distinct viruses would be surprising.

As we mentioned previously, RNA viruses have been responsible for the vast majority of recorded zoonoses in human populations (Geoghegan et al., 2017; Kreuder Johnson et al., 2015) and are often pathogenic (Ebola, SARS, Dengue, Zika, Influenza, HIV, etc.). Since the VIPs of specific viruses can be used as proxies for their ancient viral relatives, we use 2,691 RNA VIPs and 2,604 DNA VIPs as proxies for looking at sweeps left by ancient RNA viruses or by ancient DNA viruses, respectively.

We estimate the enrichment of recent sweeps at RNA VIPs by comparing RNA VIPs with all other protein coding genes far (>500kb) from RNA VIPs, including both non-VIPs and other VIPs that do not interact with RNA viruses. Similarly, we estimate the enrichment of recent sweeps at DNA VIPs by comparing DNA VIPs with all protein coding genes located far away (>500kb). In short, we test if what matters for observing a sweep enrichment is being close to RNA VIPs, or being close to DNA VIPs. We use the bootstrap test again to match confounding factors exactly the same way we did when comparing all VIPs and non-VIPs.

Counting sweeps and summing them over all the 26 populations from the 1,000 Genomes Project, we find a substantial enrichment in strong selective sweeps at RNA VIPs (Figure 2A; whole enrichment curve P<0.001). This enrichment is reminiscent of the one observed comparing all VIPs and non-VIPs (Figure 1A,B). Conversely, when we compare DNA VIPs with genes far from them, we do not observe any enrichment of strong selective sweeps (Fig 2B). We nevertheless observe some enrichment when using the rank thresholds from top 2,000 to top 500, suggesting weaker sweeps signals at DNA VIPs. These results suggest that RNA viruses exerted a more drastic selective pressure driving a larger number of strong selective events compared to DNA viruses during recent human evolution. The past 50,000 years of human evolution might thus echo the very strong skew of recorded zoonoses toward RNA viruses.

**Figure 2.**
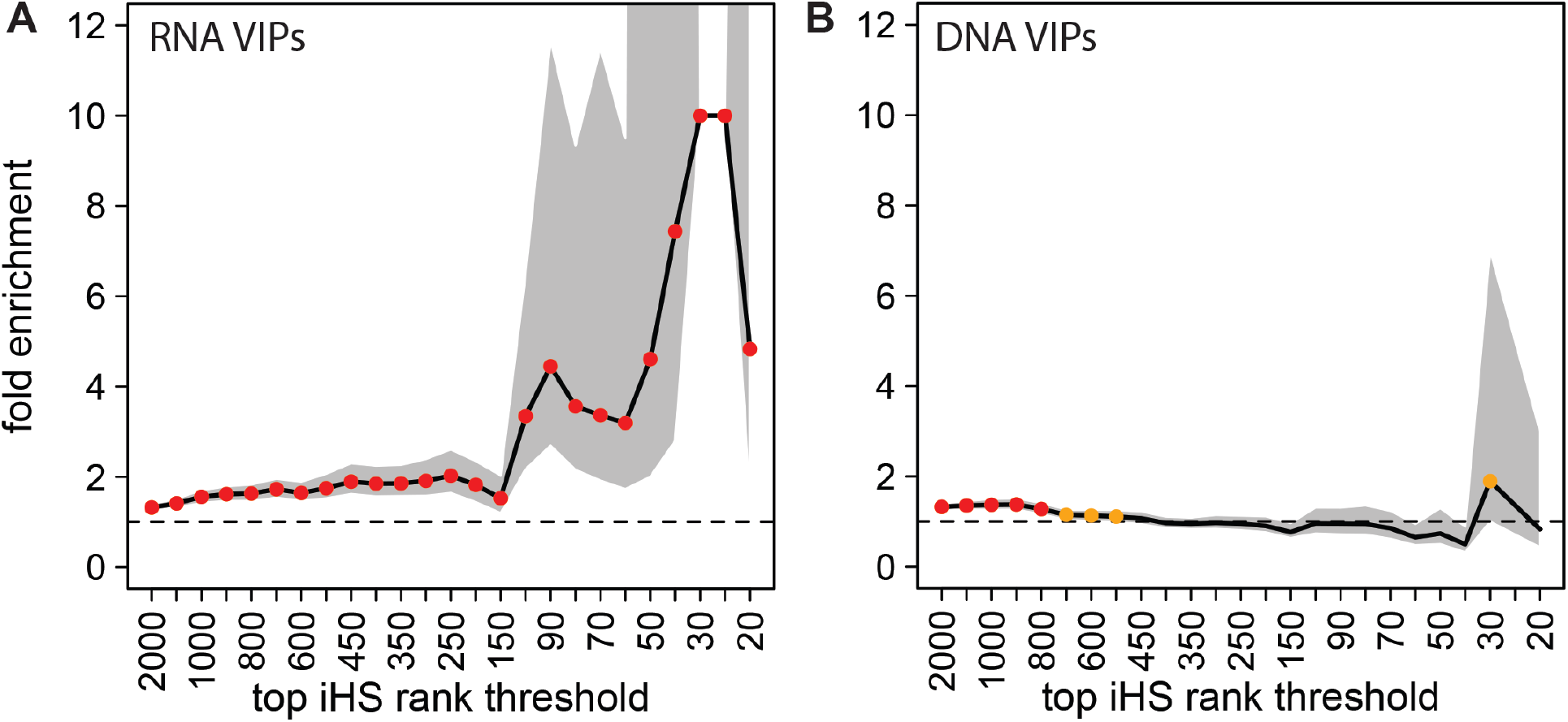
Sweep enrichment at RNA VIPs and DNA VIPs. Legend same as figure 1. A. Sweep enrichment at RNA VIPs compared to other genes far from RNA VIPs (>500kb). The enrichment exceeds tenfold at iHS top 30 and 25, and is represented at ten without confidence intervals. B. Sweep enrichment at DNA VIPs compared to other genes far from DNA VIPs (>500kb). Red dots: bootstrap test P<0.001. Orange dots: P<0.05.

### Host functions do not explain the sweep enrichment at VIPs or at RNA VIPs

Even though we control for a number of confounding factors in the previous comparisons (VIPs vs. non-VIPs, RNA VIPs vs. other genes and DNA VIPs vs. other genes), we have so far not controlled for the potential confounding effect of the host biological functions on sweep enrichments. Because the representation of specific host functions can be different in VIPs and non-VIPs, these host functions rather than interactions with viruses might explain the sweep enrichment at VIPs, specifically at RNA VIPs. Specific host functions may indeed be enriched in selective sweeps, and also over-represented in VIPs compared to non-VIPs (or RNA VIPs compared to other genes). For example, one might think of a hypothetical case where VIPs are enriched for sweeps not because they interact with viruses, but because they are enriched in cell cycle proteins, and cell cycle proteins are enriched for sweeps regardless of whether they interact with viruses or not.

To test the possibility that host functions may explain our results instead of viruses, we use functional annotations from the Gene Ontology (GO) (Gene Ontology, 2015) and first ask which GO biological annotations are enriched in VIPs compared to non-VIPs (at least 100% enrichment, lower enrichments are unlikely to explain the several fold excess of sweeps observed at VIPs). We then build a large group of genes, each gene belonging to at least one of the enriched GO annotations. We then use the bootstrap test to compare sweeps in this group with sweeps in control genes that belong to none of the VIP-enriched GO annotations (Fig. 3). If GO annotations rather than viruses explain our results, we expect the genes from the VIP-enriched GO annotations to be significantly enriched for sweeps compared to other genes. Because the confounding effect should be independent from viruses, we should be able to observe the confounding sweep enrichment using only non-VIPs, after having completely excluded VIPs from the comparison test. When we use the bootstrap test to compare non-VIPs within VIP-enriched GO annotations, and control non-VIPs, we do not find any particular sweep enrichment (whole sliding rank threshold curve P=0.27; see Methods). We do not detect any sweep enrichment when testing with GO annotations over-represented (100% or more) in RNA VIPs either (whole curve P=0.62). These tests show that the host intrinsic biological functions are unlikely to explain our results, which further supports the role of viruses, and in particular RNA viruses, in the observed patterns of recent adaptation.

**Figure 3.**
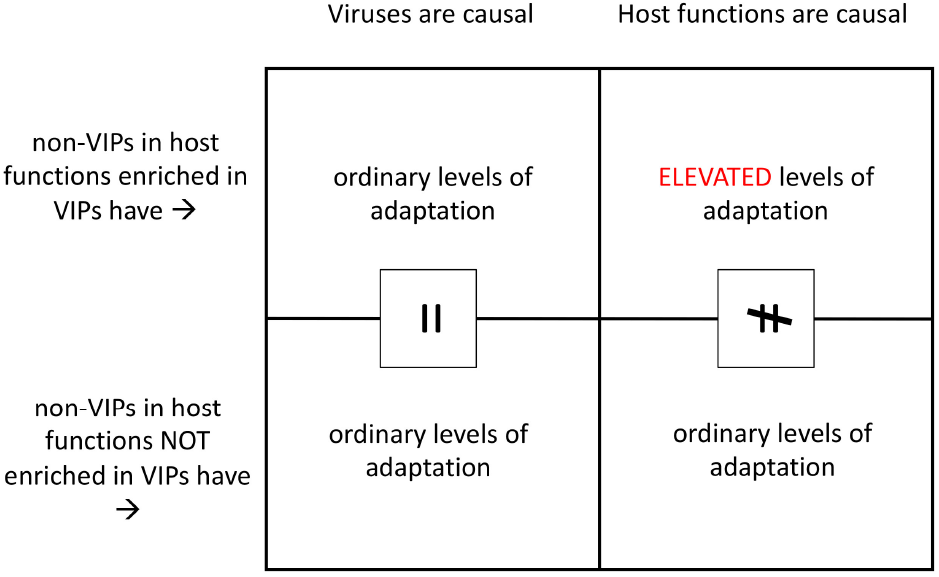
Rationale of the test for confounding host intrinsic functions. If host functions cause the enrichment in adaptation at VIPs, then these over-represented host functions in VIPs concentrate the bulk of adaptation in VIPs but also in non-VIPs. This implies that non-VIPs in those host functions should exhibit more adaptation than non-VIPs in other host functions (right side of the table). If there is no difference (left side of the table), then host functions do not confound our analysis, and viruses are likely causal.

## Discussion

We find a sharp difference in recent sweep enrichment at VIPs compared to non-VIPs, and more specifically at RNA VIPs compared to DNA VIPs worldwide. Even though the enrichment of sweeps at RNA VIPs is very unlikely to have happened by chance alone, or because of confounding factors, our results must be considered as preliminary. Because the enrichment analysis we conducted represents a correlation of sweeps signals with the location of RNA VIPs in the genome, it does not fully establish that viruses caused the sweeps. Establishing causality will require more work in the following future directions.

First, identifying the variants causal to the sweeps will help establishing plausible functional mechanisms of adaptation in response to viruses. For example, several selective sweeps might have been driven by causal non-synonymous variants localized in the contact interface between the host VIP and a viral protein, at a site in the contact interface known to be important for the physical interaction. Alternatively, the sweep causal variants might correspond to expression quantitative trait loci (eQTLs), with the selected alleles corresponding to either alleles increasing the expression of antiviral VIPs, or to alleles decreasing the expression of proviral VIPs that a virus needs to replicate. In short, the identification of causal functional alleles affecting coding or regulatory sequences in a way expected to be detrimental to viral replication will provide clear support to the causal role of viruses behind the sweep enrichment observed so far. A potential obstacle is that we specifically observe strong enrichment in large selective sweeps driven by strong recent adaptation. This may complicate the identification of the causal variants if the large sweeps include many linked variants. That said, new methods were recently published with increased power and accuracy to isolate the causal variants even in large sweeps (Akbari et al., 2018).

Second, we have so far identified a worldwide sweep enrichment at RNA VIPs, considering RNA viruses as one broad category. However, specific viruses are expected to have driven selective sweeps in specific human populations. Detecting sweep enrichments in specific human populations at VIPs that interact with specific RNA viruses may therefore further support causality, in addition to further identifying which viruses were particularly active during ancient epidemics. An important obstacle to detecting the effect of specific viruses in specific human populations is that the set of VIPs that interact with a specific virus can be substantially smaller than the set of all VIPs or RNA and DNA VIPs. For example, of the 2691 RNA VIPs, only 215 interact with Ebola virus. In order to make up for the loss of statistical power that is inevitable when using smaller samples of VIPs, we will have to use sweep detection tools beyond the single iHS statistic, with improved power and accuracy. Specific machine learning approaches that combine multiple summary statistics may offer increased performance compared to the use of a single statistic (Schrider and Kern, 2016, 2017; Sugden et al., 2018).

Third, if viruses were causal, we expect that the selective sweeps at VIPs of a specific virus would have happened around the same evolutionary time within a specific human population, which would correspond to the time of an epidemic, as opposed to being dispersed randomly across different evolutionary times with no connection to a specific epidemic event. Dating selective sweeps linked to a specific virus, with a significant convergence in individually estimated evolutionary times, may thus strengthen the evidence that viruses were causal, in addition to providing estimates of when they drove adaptation. Using combinations of summary statistics instead of only one may again provide more accuracy when dating selective events with approaches such as Approximate Bayesian Computation (Nakagome et al., 2016).

Finally, the findings that both recent selective sweeps and adaptive introgression from Neanderthals to Eurasian modern humans were dominated by RNA viruses, raises the important question of whether or not RNA viruses systematically drove more adaptation during most of human evolution, and more broadly during the evolution of other species. More work is now needed to quantify the impact of RNA viruses compared to DNA viruses at deeper evolutionary time scales in humans, using for example recent implementations of the McDonald-Kreitman test (Galtier, 2016; Tataru et al., 2017; Uricchio et al., 2019), and to quantify the impact of RNA viruses in other species both during recent and deeper evolution. It will also be interesting to see if the genes involved in recent human adaptation to RNA viruses, were also more often involved in adaptation over deeper evolutionary time scales, which would highlight genes that may be particularly important for host adaptation. Because RNA molecules from ancient RNA viruses may prove very hard to impossible to recover, our results potentially illustrate the importance of pursuing host genome-based, indirect approaches to uncover important pathogenic players in host evolution.

## Methods

### Measuring iHS and power analysis

The iHS summary statistic is computed for each variant in a genome with a minor allele frequency greater than 5%, and known derived and ancestral alleles. For our analysis we used the hapbin software (Maclean et al., 2015) to rapidly measure iHS for variants in all the 26 separate human populations represented in the 1,000 Genomes Project phase 3. We then computed the average of |iHS| (iHS can take both highly negative or positive values that both indicate adaptation) across all the variants within large 1,000kb windows. Large 1,000kb windows are more specifically sensitive to large sweeps driven by strong adaptation (see power analysis below). To assign a iHS rank to each gene, each window was centered at the genomic center of a gene, half-way between the most upstream transcription start site and the most downstream transcription end site. The gen coordinates were obtained from Ensembl v83 (ref). This window configuration avoids introducing biases related to gene length. Indeed, one can imagine an alternative way to rank genes by iHS where the value used to rank genes is the average iHS measured only for variants that overlap a gene. The problem with this is that longer genes would then be more likely to overlap high local |iHS| values just by chance compared to shorter genes, thus biasing power in favor of larger genes. For this reason, we prefer to use a constant window size.

To determine a window size that would more specifically detect strong adaptation in the human genome, we ran population simulations of incomplete selective sweeps and measured the power of iHS to detect weak or strong adaptation when using different window sizes. In particular, we used two window sizes, with small 50kb windows and much larger 1,000kb windows. We ran the population simulations using discoal (Kern and Schrider, 2016). Each simulation included 50 individuals (100 chromosomes) and represented a locus of 1.2 megabases with a uniform recombination rate of 1 cM/Mb, and a total cumulated theta of 1,800 to match the average recombination rate and diversity observed in the 1,000 Genomes Project populations. In order to measure the statistical power of iHS windows, we first had to determine the distribution of the corresponding iHS values in the neutral case without selective sweeps. This distribution was obtained running 10,000 independent neutral simulations. Note we simulated a constant population size of 10,000 since fluctuations in population size are not likely to change the difference in power between small or large windows of iHS. We simulated a large number of neutral loci to get precise power estimates even at low false positive rates. Specifically, we estimated power at a low, 0.1% false positive rate.

Figure 4 shows the power of the average |iHS| to detect a range of incomplete selective sweeps (see figure legend) using either 50kb or 1,000kb windows. From the figure it is clear that 50kb windows have good power to detect sweeps for selection intensities between 2Ns=100 and 1,000, whereas 1,000 kb windows only have good power to detect sweeps with selection intensities greater than 200 (1% selection coefficient in the human genome). Large, 1,000kb windows are therefore more appropriate to detect more specifically strong adaptation events expected from interactions with viruses. For this reason, we used 1,000kb windows for the whole analysis.

**Figure 4.**
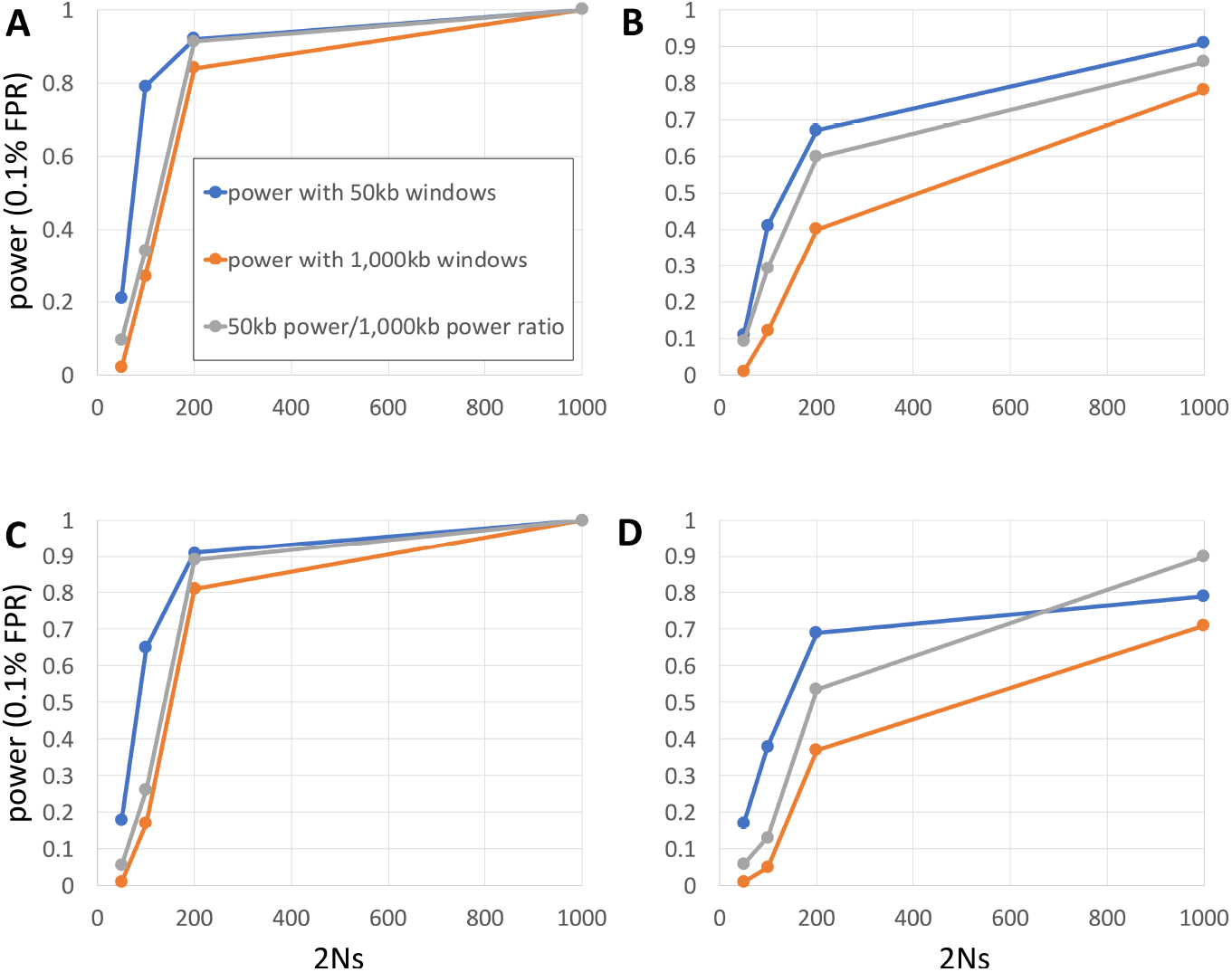
Power of iHS windows to detect various sweeps. A. Incomplete selective sweeps from a de-novo mutation that reached a 50% frequency. B. Sweep from a standing, 5% standing variant at the start of selection and that reached a 50% frequency. C. and D. same as A. and B., respectively, but for sweeps that reached a 70% instead of 50% frequency. The y axis represents the statistical power (true positive rate) at 0.1% false positive rate (FPR). The x axis represents the range of simulated selection intensities, ranging from 2Ns=50 to 2Ns=1,000 where N=10,000 in our simulations, and s is the selection coefficient. Blue curve: power with 50kb windows. Orange curve: power with 1,000kb windows. Grey curve: ratio of the power with 1,000kb windows over the power with 50kb windows.

### Testing enrichments with the boostrap test

The bootstrap test we used to match VIPs with control non-VIPs that match for multiple confounding factors has already been described extensively in a previous manuscript that the reader can refer to for more ample details (Castellano, 2019). An implementation of the bootstrap test is available at https://github.com/DavidPierreEnard, as part of a larger pipeline that also estimates the whole enrichment curve p-value (see below). In brief, the bootstrap test uses a straightforward control set-building algorithm that adds control genes to a progressively growing control set, in such a way that the growing control set has the same range of values of confounding factors as the tested set of genes of interest. For this analysis, the confounding factors that were used in the bootstrap test include the following factors likely to impact the frequency of selective sweeps:

- the density of coding sequences in 50kb windows centered on genes. Coding sequences are Ensembl v83 coding sequences.
- the density of mammalian phastCons conserved elements (Siepel et al., 2005) (in 50kb windows), downloaded from the UCSC Genome Browser (https://genome.ucsc.edu/).
- the density of regulatory elements, as measured by the density of DNASE1 hypersensitive sites (in 50kb windows) also from the UCSC Genome Browser.
- GC content (in 50kb windows).
- recombination rate from Hinch et al. (Hinch et al., 2011) in 200kb windows. We used 200kb windows instead of 50kb windows to reduce the noise in recombination rate estimates.
- average overall expression in 53 GTEx v7 tissues (Consortium, 2015) (https://www.gtexportal.org/home/). We used the log (in base 2) of RPKM values.
- expression (log base 2 of RPKM) in GTEx lymphocytes. Expression in immune tissues is likely to impact the rate of sweeps.
- expression (log base 2 of RPKM) in GTEx testis. Expression in testis is also likely to impact the rate of sweeps.
- the number of protein-protein interactions (PPIs) in the human protein interaction network (Luisi et al., 2015). The number of PPIs has been shown to influence the rate of sweeps (Luisi et al., 2015). We use the log (base 2) of the number of PPIs.
- the proportion of immune genes. The control sets have the same proportion of immune genes as VIPs, as annotated by the GO terms GO:0002376 (immune system process), GO:0006952 (defense response) and GO:0006955 (immune response), to avoid confusing the effect of viruses with broader immune effects.
- the proportion of genes that interact with bacteria according to the Intact database as of April 2018 (https://www.ebi.ac.uk/intact/). Note that we also attempted to match the proportion of genes that interact with Plasmodium (Ebel et al., 2017), but VIPs and non-VIPs had such different numbers of Plasmodium-Interacting genes that the bootstrap test failed to match them. However, we were able to use the bootstrap test to verify that plasmodium-interacting genes do not have more iHS sweeps than other genes (bootstrap test P>0.05 for all thresholds from top 2,000 to top 20).

It is important to note that we match many (11) confounding factors, which inevitably restricts the number of non-VIPs that can be used as controls. Smaller numbers of control non-VIPs increase the false positive rate of the bootstrap test (see below). However, the False Discovery Rate analysis strategy that we use to test the significance of the whole curve enrichment fully takes this limitation into account (see below). Note also that in order to limit the number of false positive sweeps due to low recombination regions included in the bootstrap test, we did not use genes with recombination rates estimated to be lower than 0.2 cM/Mb (Hinch et al., 2011).

### Testing the significance of the whole enrichment curve

In addition to estimating a p-value for each iHS rank threshold from top 2,000 to top 20 with the bootstrap test, we also estimated how significantly the whole enrichment curve stands above null expectations. We do this for multiple reasons, some of which already mentioned above. First, the different thresholds across the whole curve are not independent from each other. As a consequence, an excess of significant p-values at multiple thresholds may reflect a correlation of false positives due to the dependence between thresholds. Second and most importantly, it is very likely that we used the bootstrap test in conditions that make it non-nominal (meaning a non-uniform distribution of p-values under the null hypothesis of no enrichment). Indeed, as we mentioned in the results, we had to exclude many non-VIPs too close to VIPs (<500kb) to avoid counting VIP sweeps as non-VIP sweeps, and vice versa. We also used only non-VIPs that could match VIPs for a large number of confounding factors. The issue then is that we only had a limited number of non-VIPs that were far enough from VIPs and that could be used as controls. For example, for the VIPs vs. non-VIPs test, we could use only 1,704 control non-VIPs. A smaller set of controls means that in the bootstrap test, the same control non-VIP has to be re-sampled more times (it is a bootstrap), thus resulting in a smaller set of control non-VIPs. Smaller samples of control non-VIPs are inevitably related to (i) a higher variance of the control null distribution, and (ii) also to a higher variance of the overall average of the null distribution. Bias (i) can result in a higher rate of false negatives with a decreased power to detect an significant sweep enrichment, but more alarmingly bias (ii) can on the contrary inflate the rate of false positive bootstrap tests if the average of the small control sets is far from what the average would be for ideal, very large control sets.

This is a very serious limitation of our analysis, but fortunately there is a very simple, yet computationally intensive remedy. Since we know that the bootstrap test is likely not nominal, we can estimate the true False Discovery Rate associated with the whole approach, by re-running the entire analysis pipeline many times on randomized genomes where the iHS ranks have been swapped randomly between genes. We specifically use 10,000 random genomes. We can then estimate how significant the real enrichment curve from top 2,000 to top 20 is compared to the same curved measured on the 10,000 random genomes. As a statistic to estimate the significance of the whole curve, we use the difference of the observed number of VIPs, minus the expected number according to controls, and we sum this difference over all rank thresholds from top 2,000 to top 20 (Figure 1). We then compare the real value of the statistic for the real genome, with the distribution of 10,000 random values obtained from the 10,000 randomized genomes. For each randomized genome, we use exactly the same VIPs and control non-VIPs as we did when testing the real genome, but these genes are now associated with randomly swapped iHS ranks. The sample size of the control sets of non-VIPs is exactly the same using random genomes as when using the real genome, meaning that the results of the boostrap test reproduce exactly the same biases for the random genomes as when testing the real genome. The p-value obtained for the whole rank thresholds enrichment curve after 10,000 randomized genomes is thus an unbiased, nominal p-value that matches the actual False Discovery Rate.

It is however very important to note that it is true that the randomized genomes provide an unbiased, whole curve test, only because we do not randomize genomes in a completely random fashion. If we just swapped randomly iHS ranks between genes, we would lose the very important property of the real genome that genes with top iHS ranks are likely to be neighbors and to form clusters. Indeed, selective sweeps, and especially strong large selective sweeps, can overlap with multiple genes in the human genome, thus creating clusters of neighboring high-ranking iHS genes. In our case, it was therefore crucial to randomize iHS ranks between genes in a way that conserved the same exact clustering structure between top iHS rank genes. To achieve this, we cut the genes ordered as they are across human chromosomes in 100 blocks of contiguous genes, and then randomly shuffled these blocks. The size of the blocks is much larger than the size of even very large sweeps of multiple megabases, thus ensuring that the clustering structure of the top iHS rank genes is preserved within the blocks. Randomly swapping the blocks still results in randomly swapping iHS ranks between genes within the blocks. Because the randomized genomes obtained this way preserve the clustering of iHS signals, we can still measure enrichments using the number of genes, instead of having to count the number of sweeps. Note that the entire pipeline to get the whole enrichment curve p-value is available at https://github.com/DavidPierreEnard with a user manual, together with the bootstrap test.

### Testing the involvement of host intrinsic functions with GO categories

To test whether host intrinsic functions can explain the sweep enrichment at VIPs and particularly at RNA VIPs, we measured enrichment for sweeps among genes that belong to GO functions that are over-represented in VIPs. Specifically, we tested whether genes that belong to over-represented functions in VIPs are enriched for sweeps completely independently of whether they interact with viruses or not, by completely excluding VIPs and genes at less than 500kb from VIPs (and more likely to overlap VIP sweeps) from the analysis. Thus, we used the bootstrap test to compare iHS sweeps in 718 non-VIPs more than 500kb from VIPs that belong to over-represented GO functions in VIPs (GO functions had to be found in 50 VIPs or more, 100% over-represented or more), with sweeps in 423 non-VIPs more than 500kb from VIPs that do not belong to over-represented GO functions in VIPs. The 718 non-VIPs with over-represented GO functions are themselves at least 500kb from non-VIPs with no over-represented GO function to avoid the issue of sweeps overlapping the two categories. For RNA VIPs, we used 1,726 nonRNA-VIPs within over-represented GO functions, and 357 nonRNA-VIPs with no over-represented GO function. We ran the bootstrap test testing for a deficit of sweeps in the 423 non-VIPs out of over-represented GO functions using the larger number of non-VIPs with at least an over-represented function (the larger the pool of controls compared to the tested set, the better). We also flipped the two groups when testing GO functions for RNA VIPs for the same reason that it is always better to have the larger group being the control group in the bootstrap test. We estimated the significance of the bootstrap test by measuring the whole enrichment curve p-value as already described above. We used the same confounding factors to match during the bootstrap test as we previously did to match VIPs with control non-VIPs.

### Functional relevance of VIPs

To estimate the functional relevance of VIPs for the viral replication cycle, we counted how many manually curated VIPs from the first 200 rows in Table S1 had clear reported proviral or antiviral effects when their function was experimentally perturbed either via expression perturbation or via protein function perturbation. The experimental perturbations had to be reported in the publications listed in Table S1. We found that 66% of the interactions with VIPs had clear functional consequences for the viral replication cycle.

